# Doublecortin-like kinase 1 (DCLK1) Facilitates Dendritic Spine Growth of Pyramidal Neurons in Mouse Prefrontal Cortex

**DOI:** 10.1101/2022.05.13.491846

**Authors:** Kelsey E. Murphy, Erin Y. Zhang, Elliott V. Wyatt, Justin E. Sperringer, Bryce W. Duncan, Patricia F. Maness

**Affiliations:** Department of Biochemistry and Biophysics, and Carolina Institute of Developmental Disabilities, University of North Carolina School of Medicine at Chapel Hill

**Keywords:** Doublecortin-like kinase, dendritic spines, cortical pyramidal neurons, NrCAM, Ankyrin, mouse models

## Abstract

The L1 cell adhesion molecule NrCAM (Neuron-glia related cell adhesion molecule) functions as a co-receptor for secreted class 3 Semaphorins to prune subpopulations of dendritic spines on apical dendrites of pyramidal neurons in the developing mouse neocortex. The developing spine cytoskeleton is enriched in actin filaments but a small number of microtubules have been shown to enter the spine apparently trafficking vesicles to the membrane. Doublecortin-like kinase 1 (DCLK1) is a member of the Doublecortin (DCX) family of microtubule-binding proteins with serine/threonine kinase activity. To determine if DCLK1 plays a role in spine remodeling, we generated a tamoxifen-inducible mouse line (Nex1Cre-ERT2: DCLK1^flox/flox^ : RCE) to delete microtubule binding isoforms of DCLK1 from pyramidal neurons during postnatal stages of spine development. Homozygous DCLK1 conditional mutant mice exhibited decreased spine density on apical dendrites of pyramidal neurons in the prefrontal cortex (layer 2/3). Mature mushroom spines were selectively decreased upon DCLK1 deletion but dendritic arborization was unaltered. Mutagenesis and binding studies revealed that DCLK1 bound NrCAM at the conserved FIGQY^1231^ motif in the NrCAM cytoplasmic domain, a known interaction site for the actin-spectrin adaptor Ankyrin. These findings demonstrate that DCLK1 facilitates spine growth and maturation on cortical pyramidal neurons in the mouse prefrontal cortex potentially through microtubule and NrCAM interactions.

## Introduction

Dendritic spine remodeling in developing pyramidal neurons is important for establishing an appropriate level of excitatory connections in the neocortex. The L1 family cell adhesion molecules Neuron-glia related protein (NrCAM) and Close Homolog of L1 (CHL1) mediate selective pruning of dendritic spine subpopulations in the mouse neocortex in response to class 3 secreted Semaphorins (Sema3s) through an activity dependent mechanism (Tran et al., 2009;Mohan et al., 2019a;Mohan et al., 2019b). Heterotrimeric receptors comprising L1-CAMs, Neuropilins (Npn1-2), and PlexinAs (PlexA1-4) transduce Sema3 signals to achieve selective spine pruning (Duncan et al., 2021a;Duncan et al., 2021b). All L1-CAMs share a conserved cytoplasmic domain containing the motif FIGQY (FIGAY in CHL1) that binds the actin-spectrin adaptor protein Ankyrin when the tyrosine (Y) residue is not phosphorylated (Bennett and Healy, 2009).

Doublecortin-Like Kinase 1 (DCLK), a member of the doublecortin (DCX) family, is a microtubule-associated protein with serine/threonine kinase activity, important for microtubule bundling and linkage to F-actin (Friocourt et al., 2007;Yap et al., 2012;Yap and Winckler, 2015;Lipka et al., 2016). DCLK1 on human chromosome 13 is associated with schizophrenia (Havik et al., 2012), whereas X chromosome mutations in DCX result in lissencephaly (smooth brain) and a double-layered cortex, producing intellectual disability (des Portes et al., 1998). Recently, DCLK1 was reported to regulate the levels of α-Synuclein, a protein that induces neurotoxicity at increased levels, and may drive a genetic form of Parkinson’s Disease (Vazquez-Velez et al., 2020). As shown in Fig.1A, DCX contains two microtubule binding domains and a serine/proline rich PEST (proline, glutamate, serine, threonine) sequence. DCLK1 (82 kDa) also has two microtubule binding domains and the PEST sequence, as well as a carboxyl terminal serine/threonine protein kinase domain. Alternative splicing of the DCLK1 gene generates a DCX-like isoform of 40 kDa, lacking the kinase domain, while alternative promoter usage generates a short isoform, DCLK1-S (42 kDa), lacking the microtubule binding domains. Substrates of DCLK1 are poorly defined but include DCLK1 itself, which inhibits microtubule bundling and polymerization (Patel et al., 2016), and microtubule associated protein 7 domain 1 (MAP7D1), which promotes axon elongation (Koizumi et al., 2017). Several lines of evidence suggest that DCLK1 may regulate spine morphogenesis. (a) DCLK1 is present in the somatodendritic region of neurons (Shin et al., 2013) and transports KIF1-mediated cargo of dense core vesicles along microtubules to increase dendrite growth (Lipka et al., 2016). (b) Microtubules have been shown to invade spines for protein and vesicle transport via kinesin to facilitate synaptic plasticity (McVicker et al., 2016;Dent, 2017). (c) DCLK1 knockdown by shRNA in embryonic neural progenitors decreases dendritic arborization and alters spine morphology (Shin et al., 2013). Although global knockout of DCLK1 in mice showed intact neuronal migration, cortical thickness, and lamination, subtle phenotypic changes in neurons, including spine density, were not examined (Deuel et al., 2006;Shin et al., 2013).

To investigate a function for DCLK1 in postnatal cortical pyramidal neurons, we generated an inducible, conditional mouse line (Nex1Cre-ERT2: DCLK1^flox/flox^ : RCE) targeting the principal microtubule binding isoforms. Tamoxifen induction of the CreERT2 recombinase under control of the Nex1 promoter achieves cell-specific targeting of postmitotic cortical and hippocampal pyramidal neurons with no detectable targeting of interneurons, oligodendroglia, astrocytes, or non-neural cells (Agarwal et al., 2012). By introducing tamoxifen at early postnatal stages gene excision can be restricted to postmigratory pyramidal neurons, circumventing effects on migration and cortical positioning. This induction regimen was applied during the active period of spine remodeling (P10-P13) in Nex1Cre-ERT2: DCLK1^flox/flox^ : RCE mice enabling examination of DCLK1 function in dendritic spine regulation in postmitotic pyramidal neurons.

Homozygous recombination of DCLK1 in Nex1Cre-ERT2: DCLK1^flox/flox^ : RCE mice during postnatal spine remodeling decreased DCLK1 expression, spine density and mature morphology on apical dendrites of pyramidal neurons in the mouse prefrontal cortex. In addition, DCLK1 bound the neural adhesion molecule NrCAM, a known regulator of spine density (Demyanenko et al., 2014;Mohan et al., 2019a), and specifically targeted the Ankyrin interaction motif (FIGQY) in the NrCAM cytoplasmic domain. Generation of an inducible mouse model for DCLK1 in pyramidal neurons, and identification of a novel role for DCLK1 in dendritic spine regulation contributes to our understanding of postnatal brain development, and may shed light on pathological mechanisms associated with its dysfunction.

## Experimental Procedures

### Generation and tamoxifen induction of DCLK1 conditional mutant mice

A conditional, inducible DCLK1 mouse line (Nex1Cre-ERT2: DCLK1^flox/flox^ : RCE) was generated, in which DCLK1 can be deleted in pyramidal neurons of the neocortex and hippocampus under control of the Nex1Cre-ERT2 promoter by tamoxifen treatment at specific times in development. The Nex1 promoter drives the tamoxifen-inducible Cre-ERT2 recombinase only in post-mitotic pyramidal neurons (Agarwal et al., 2012).

As shown in Fig. 1B, Nex1Cre-ERT2 mice (originally from Amit Agarwal) were first intercrossed with RCE: loxP mice (from Gordon Fishell), in which expression of Cre causes recombination of a “floxed stop cassette” allowing EGFP expression (Sousa et al., 2009). Nex1Cre-ERT2: RCE mice were then intercrossed with DCLK1^tm1.2jgg^ mice containing floxed exon 3, the second coding exon, on a mixed genetic background of C57/Bl6, 129 and Black Swiss (Jackson Laboratory strain 013170; (Koizumi et al., 2006)), Joseph Gleeson, donating investigator). Cre-mediated recombination has been shown to delete exon 3, eliminating expression of full length DCLK1 (82K) and the alternatively spliced DCX-like variant (40 kDa) lacking the kinase domain (Koizumi et al., 2006). Subsequent breeding produced Nex1Cre-ERT2: RCE mice with DCLK1^flox/flox^, DCLK1^flox/+^, and DCLK1^+/+^(wild type, WT) genotypes in expected ratios. Each allele was genotyped by PCR from tail DNA (Fig. 1D). For *in vivo* induction, tamoxifen (Sigma-Aldrich, #10540-29-1) was dissolved (10 mg/ml) in sunflower seed oil (Sigma, S5007), and administered by intraperitoneal injection at 100 mg/kg body weight every 24 hours for 4 consecutive days. Balanced sexes of male and female mice were used in the present studies. Mice were maintained according to policies of the University of North Carolina Institutional Animal Care and Use Committee (IACUC; AAALAC Institutional Number: #329; ID# 18-073, 21-039) in accordance with NIH guidelines. Genotyping was performed by PCR.

**Figure 1:**
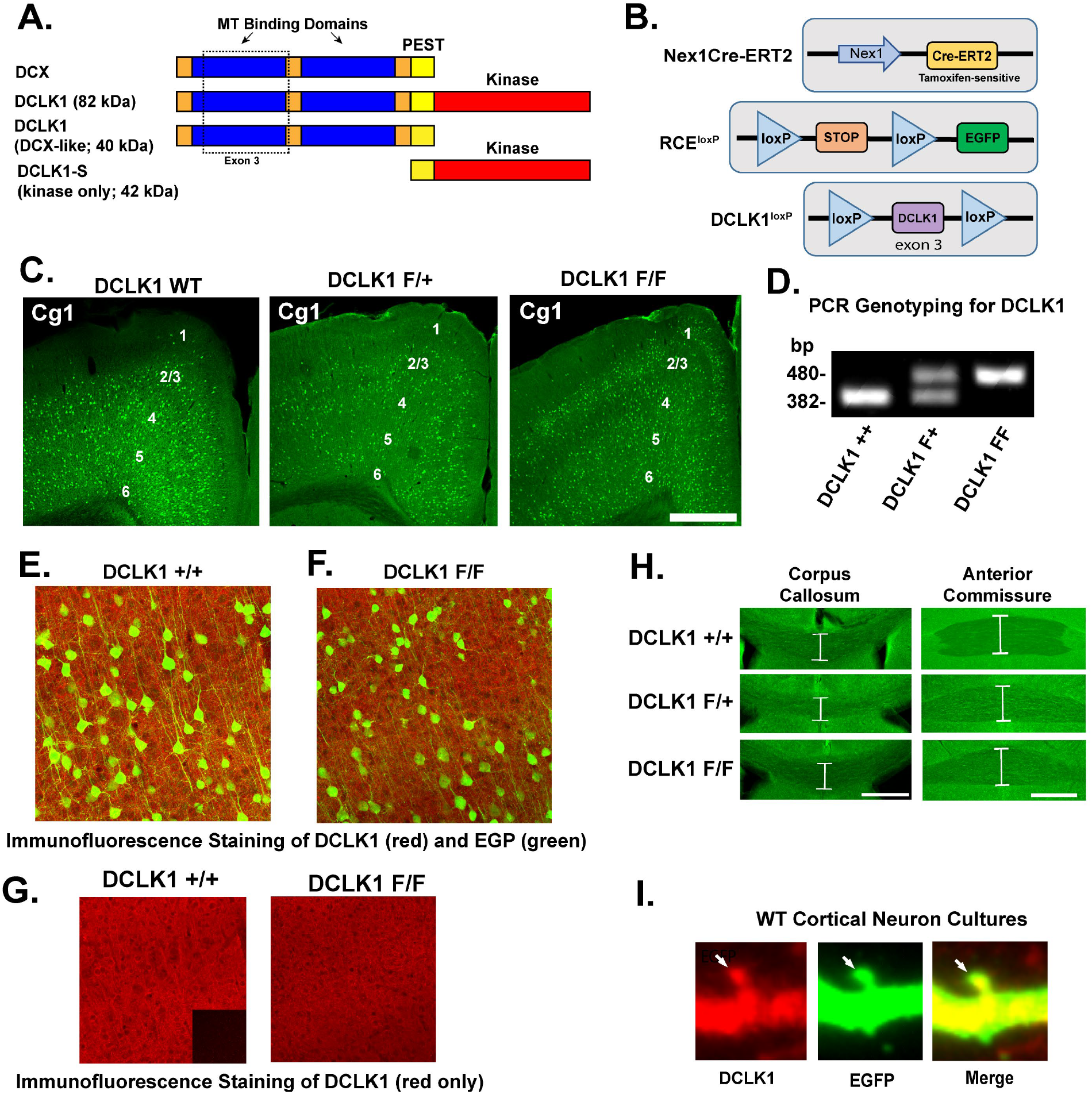
Conditional deletion of DCLK1 in postnatal pyramidal neurons of Nex1Cre-ERT2: DCLK1 F/F: RCE mice. (A) DCX and DCLK1 isoforms and domain structure. MT, microtubule; PEST sequence; serine/threonine kinase domain. (B)Cre-Lox diagram for recombination in Nex1Cre-ERT2: DCLK1 F/F: RCE mice. (C)Recombination in DCLK1 +/+, F/+ and F/F cingulate cortex (Cg1; P24) indicated by EGFP expression after tamoxifen induction at P10-13. Scale bar = 500 µm. (D)PCR genotyping was used to identify exon 3 excision in DCLK1 DNA extracted from DCLK1 +/+, F/+ and F/F mice, as indicated by the size of diagnostic bands on an agarose gel. (E-F) Immunofluorescence staining of DCLK1 protein (red) in EGFP-expressing pyramidal neurons (green) in Cg1 layer 2/3 of DCLK1 +/+ and DCLK1 F/F mice after tamoxifen induction, shown in merged images. (G) Single channel DCLK1 immunofluorescence staining (red) in DCLK1 +/+ and DCLK1 F/F Cg1 after tamoxifen induction as in E,F. A secondary antibody control without primary antibodies is shown as an inset. (H)The width of the corpus callosum (CC) and anterior commissure (AC) (white brackets) was measured at the midline in serial coronal brain sections in DCLK1 +/+, F/+, and F/F mice expressing EGFP. DCLK1 F/F mice (P19-34) induced at P10-P13 showed no change in width of these axon tracts compared to DCLK1 +/+ or DCLK1 F/+ mice (see Results section, using 1 factor ANOVA with Tukey’s post hoc tests; p< 0.05; P19-34). Scale bar = 300 µm. (I)Representative images of apical dendritic spines in cortical neuron cultures, immunostained for DCLK1 (red) and EGFP (green), and a merged image. WT cortical neuronal cultures at DIV14 were transfected with pCAG-IRES-EGFP, immunostained for DCLK1 and EGFP, and imaged confocally. DCLK1 immunoreactivity was present on dendritic shafts and spines as shown in the representative image. Arrows indicate a mushroom-shaped spine on an apical dendrite.

### Immunoreagents

Rabbit polyclonal antibodies against the carboxyl-terminal region (residues 700 to C-terminus) of mouse DCLK1 (Abcam, # ab31704) recognize full length DCLK1 and the short isoform DCLK1-S. These antibodies have been knockout validated. Polyclonal antibodies against NrCAM (Abcam #24344) or R&D Systems (Abcam #8538) and GFP (Abcam #13970) were also used. Non-immune rabbit IgG (NIg), HRP- and AlexaFluor (488 and 555) conjugated secondary antibodies were from Jackson Immunoresearch.

### Analysis of spine density, morphology, and dendritic branching

Mice (P19-P34) were anesthetized with 2.5% Avertin, perfused transcardially with 4% paraformaldehyde (PFA)/PBS and processed for staining as described (Demyanenko et al., 1999). Brains were postfixed in 4% PFA overnight at 4°C, followed by 0.02% PBS-azide, then sectioned coronally on a vibratome (60 µm) and mounted on glass slides. Sections were permeabilized in 0.3% Triton X-100 and blocked in 10% normal donkey serum in PBS for 3 hours at room temperature. Sections were incubated with chicken anti-GFP (1:250) and anti-DCLK1 antibodies (1:250) for 48 hours at 4°C. After washing, sections were incubated with anti-chicken Alexa Fluor-488 or anti-rabbit Alexa Fluor-555 secondary antibodies (1:250) for 2 hours before mounting with Prolong Glass (Thermofisher). Confocal z-stacks were obtained by imaging on a Zeiss LSM 700 microscope in the UNC Microscopy Services Laboratory with an EC Plan Neofluar 40x objective with a 1.3 oil lens. Images were acquired using a pinhole size of 1 AU. Zoom was adjusted to obtain pixel sizes of 0.13-0.14 µm.

Spine density of layer 2/3 pyramidal neurons in the prefrontal cortex (primary cingulate area) was quantified using Neurolucida software (MBF Bioscience) as described (Demyanenko et al., 2014;Mohan et al., 2019a;Mohan et al., 2019b). Briefly, spines were traced and quantified blind to observer on 30 µm segments of the first branch of apical or basal dendrites from confocal z-stack images after deconvolution. For image deconvolution and 3D reconstructions, dendritic z-stacks were deconvolved using AutoQuant 3 software (Media Cybernetics) with default deconvolution settings in Imaris (Bitplane). Mean spine number per 10 µm of dendritic length (density) was calculated. Approximately 40-50 neurons/mouse were analyzed (4-6 mice/genotype; P19-P34). Mean spine densities/10 μM ± SEM were compared across genotypes by two-factor ANOVA and Tukey’s posthoc testing for multiple comparisons, with significance set at p<0.05.

Spine morphologies were scored on 3D reconstructed images as mature (mushroom) or immature (stubby or thin/filopodial) on apical dendrites as defined (Peters and Kaiserman-Abramof, 1970;Peters and Harriman, 1990) and reported previously (Mohan et al., 2019a). To compare spine morphologies of DCLK1 +/+ (n=3) and DCLK1 F/F (n=4) mice, each spine morphological type per unit length was cpimted and the percent calculated. Mean percentages of mature (mushroom) spines and immature (thin and stubby) spines per genotype were compared for statistical significance by the t-test (2-tailed, unequal variance, p < 0.05). The number of neurons/genotype analyzed for spine morphology was 18 for DCLK1 +/+ and 19 for DCLK1 F/F.

Dendritic arborization was measured by Sholl analysis in pyramidal neurons of the cingulate cortex in DCLK1 +/+ and DCLK1 F/F mice (P19-P34, n= 3 mice/genotype) induced at P10-P13. The number of processes crossing concentric rings centered on the soma of EGFP-labeled pyramidal neurons (approximately 15 neurons/genotype) was scored in confocal z-stacks (20x). The center was defined as the middle of the cell body at a soma detector sensitivity of 1.5 µm, and the automatic tracing mode of Neurolucida was used to seed and trace dendritic arbors. Images in DAT format were subjected to Sholl analysis using Neurolucida Explorer with a starting radius of 10 µm and radial increments of 10 µm ending at 150 µm. Sholl data were compared using paired t-tests for differences in the mean number of crossings at each distance from the soma (significance set at p< 0.05).

For estimating the size of major axonal tracts (corpus callosum and anterior commissure) in DCLK1 +/+, F/+, and F/F mice (3-5 mice/genotype; P19-P34), serial coronal brain sections were made, imaged on the confocal microscope (5x), and analyzed in FIJI. The dorsoventral width of each tract was measured at the midline of each section, and the mean width calculated. Means were compared by 1 factor ANOVA with Tukey’s posthoc comparisons (p < 0.05).

### Immunoprecipitation and immunoblotting

Protein-protein interactions were assessed by co-immunoprecipitation and immunoblotted from transfected HEK293T cells, mouse forebrain, and synaptoneurosomes. HEK293T cells were grown in DMEM, gentamicin, kanamycin,10% FBS in a humidified incubator with 5% CO2. Cells were seeded at 2 × 10^6^ cells/100mm dish the day before transfection. Plasmids containing WT or mutant NrCAM were cotransfected with either Ankyrin B or DCLK1 at a 2:1 molar ratio with Lipofectamine 2000 (ThermoFisher, #11668) in Opti-MEM. Media was changed to complete DMEM after 18 h, and cells were lysed and collected 48 h post-transfection. Cells were harvested in Triton lysis buffer (20 mM Tris, pH 7.0, 150 mM NaCl, 200 μM Na3VO4, 1 mM EGTA, 1% Triton X-100, 10 mM NaF, 1x Protease Inhibitor Cocktail Sigma Aldrich #P8340). For preparation of mouse forebrain lysates, were dissociated and subjected to Dounce homogenization for 20 strokes in RIPA buffer. Homogenates were incubated for 15 minutes on ice, then centrifuged at 16,000 x g for 10 minutes. The supernatant was retained, and protein concentration determined by BCA.

Synaptoneurosomes were isolated as described (Villasana et al., 2006). WT mice (P32) were anesthetized, decapitated and cortices were isolated. Following Dounce homogenization in Triton lysis buffer, homogenates were sonicated, filtered, and centrifuged at 1000 x g for 10 minutes at 4°C. Pellets were resuspended in Triton lysis buffer, nutated, and centrifuged at 16,000 x g for 10 minutes at 4°C. Supernatants were retained as the synaptoneurosome fraction, protein concentration was determined using BCA. Postsynaptic density protein 95 (PSD95) enrichment in synaptoneurosomes over homogenate was verified by Western blotting.

For immunoprecipitation, lysates of mouse forebrain (1 mg) or HEK293T cells (0.5 mg) were precleared for 30 minutes at 4°C using Protein A/G Sepharose beads (ThermoFisher). Precleared lysates (equal amounts of protein) were incubated with 3 µg rabbit polyclonal antibody to NrCAM (Abcam #24344) or NIg for 2 h r on ice. Protein A/G Sepharose beads were added for an additional 30 min with nutation at 4°C before washing with RIPA buffer or Triton lysis buffer (synaptoneurosomes). Beads were washed 4 times, then immunoprecipitated proteins were eluted from the beads by boiling in SDS-PAGE sample buffer. Samples (50 µg) were subjected to SDS-PAGE (6%) and transferred to nitrocellulose. Membranes were blocked in TBST containing 5% nonfat dried milk and incubated overnight with primary antibodies (1:1000), washed, and incubated with HRP-secondary antibodies (1:5000) for 1 h. Antibodies were diluted in 5% milk/Tris buffered saline/0.1% Tween-20 (TBST). Blots were developed using Western Bright ECL Substrate (Advansta) and exposed to film for times yielding a linear response of signal. Membranes were stripped and reprobed with rabbit anti-NrCAM antibodies (R&D Systems AF8538). Bands were quantified by densitometric scanning using FIJI. The ratio of DCLK1 to NrCAM was determined. Normalized ratios from 3 replicate experiments for each mutant were averaged and reported ± SEM.

## Results

### Characterization of Nex1Cre-DCLK1 mutant mice

To identify functional effects of postnatal DCLK1 deletion on dendritic spine development in cortical pyramidal neurons, we generated a tamoxifen-inducible mouse line, Nex1Cre-ERT2: DCLK1^flox/flox^ : RCE (Fig.1B). In these mice, floxed exon 3 in the DCLK1 gene was excised by tamoxifen-induced loxP recombination under control of the Nex1 promoter, which is active in postmitotic pyramidal neurons (Agarwal et al., 2012;Mohan et al., 2019a). This same excision in EIIa-Cre: DCLK1 F/F mice has been shown to eliminate the DCLK1 isoforms with microtubule binding domains: full length DCLK1 (82 kDa) and DCX-like splice variant (40 kDa) (Koizumi et al., 2006). The kinase-only short isoform (DCLK1-S; 42 kDa) generated from alternative promoter usage is not deleted (Koizumi et al., 2006). DCLK1-S, also called CPG16 (Nedivi et al., 1993), is expressed exclusively in adult brain and at much lower levels than the full length isoform. Recombination was reported by EGFP expression from RCE: loxP induced upon Cre-ERT2 activation (Sousa et al., 2009). Mice were injected with tamoxifen in the juvenile period (P10-P13), when cortical layers 2-4 have mostly formed, in order to delete the microtubule binding isoforms of DCLK1 during the most active stages of spine formation and remodeling (Culotta and Penzes, 2020). To achieve recombination, mice were given daily tamoxifen injections intraperitoneally from P10 to P13 as described (Agarwal et al., 2012;Mohan et al., 2019a), resulting in permanent loss of the targeted DCLK1 isoforms. Use of the Nex1Cre-ERT2 promoter avoided deleting DCLK1 in immature progenitors, unlike Nestin or EIIa promoters. Analysis focused on pyramidal neurons in the prefrontal cortex, because of its importance in social and cognitive circuits that may be altered in DCLK1-linked neurological disease (Yizhar, 2012;Kroon et al., 2019).

Tamoxifen induction at P10-P13 resulted in expression of EGFP, indicating recombination, in pyramidal neurons throughout layers 2-6 in the primary cingulate area (Cg1) of the medial prefrontal cortex of Nex1Cre-ERT2: DCLK+/+: RCE (“WT”, termed DCLK1 +/+), heterozygous Nex1Cre-ERT2: DCLK1*flox/+*: RCE (termed F/+), and homozygous Nex1Cre-ERT2: DCLK1^*flox/flox*^: RCE mice (termed F/F) (Fig. 1C). Mice were genotyped by PCR (Fig.1D). We focused our analysis on mice at P19-P34, a juvenile to adolescent stage defined by (Laviola et al., 2003). Prior to and overlapping with this time frame, overproduced dendritic spines are pruned to appropriate levels. Spine turnover decreases substantially from P19-P34 as mature circuits are stabilized (Trachtenberg et al., 2002;Holtmaat A. J. et al., 2005).

Expression of DCLK1 protein in layer 2/3 of Cg1 was decreased in DCLK1 F/F compared to DCLK1 +/+ mice, as evident from DCLK1 immunofluorescence (red) in EGFP-labeled cell bodies and neuropil in confocal images (Fig. 1 E,F). Quantitation of DCLK1 fluorescence pixel density in the DCLK1 F/F Cg1 (Fig. 1G) was significantly reduced (55.3 ± 0.5 pixel intensity/unit area) compared to DCLK+/+ Cg1 (72.6 ± 0.5, t-test, *p<0.0001). Antibodies used for immunostaining recognized the DCLK1 carboxyl terminus, which is shared by DCLK1 (82 kDa) and DCLK1-S (42K). Residual staining may be due to DCLK-S, and to any non-recombined pyramidal neurons or other cell types. DCLK1 antibodies were also used for immunostaining of dissociated cortical neurons in culture at DIV14 expressing EGFP. DCLK1 immunolabeling was evident on dendritic shafts and spines, as shown in Fig. 1 I.

In homozygous DCLK1 F/F or heterozygous DCLK1 F/+ mice, tamoxifen induction at P10-P13 did not cause gross neuroanatomical defects or alterations in the size of large axonal tracts, in particular the corpus callosum and anterior commissure (Fig. 1H). There was no significant decrease in the width of the corpus callosum at the midline (DCLK1 +/+ (231 µm ± 39, n= 3 mice); DCLK1 F/+ (238 µm ± 24, n = 5); DCLK1 F/F (241 µm ± 38, n= 5) or that of the anterior commissure (DCLK1 +/+ (454 µm ± 13); DCLK1 F/+ (444 µm ± 35); DCLK1 F/F (424 µm ± 37) (1 factor ANOVA with Tukey’s post hoc testing; p< 0.05; P19-34). Cortical thickness and lamination, and size of the hippocampus and brain ventricles appeared unaltered in the mutant mice, and other neuroanatomical abnormalities were not apparent, in agreement with the phenotype of DCLK1 global knockout mice (Deuel et al., 2006).

### Postnatal deletion of DCLK1 from cortical pyramidal neurons decreases spine density and alters spine morphology

To investigate an *in vivo* role for DCLK1 in regulating spine density in the prefrontal cortex, recombination was induced in DCLK1 +/+, F/+ and F/F mice at P10-P13, and spine density was analyzed on EGFP-positive apical dendrites of pyramidal neurons in layer 2/3 of Cg1 at P19-P34 (Fig. 2A). Apical dendrites were analyzed because spines on apical but not basal dendrites are selectively regulated by L1-CAMs (Duncan et al., 2021b), which may interact with DCLK1. Spine density was significantly decreased on apical dendrites of cortical pyramidal neurons in DCLK1 F/F compared to DCLK1 +/+ mice (*p<0.0001, 1-factor ANOVA with Tukey’s posthoc comparisons) (Fig. 2B). DCLK1 F/+ mice also exhibited significantly decreased spine density compared to DCLK1 +/+ mice (*p = 0.002) or to DCLK1 F/F mice (*p = 0.011) (Fig. 2B). In summary, the findings suggested that DCLK1 is required to facilitate spinogenesis or maintenance in postnatal pyramidal neurons in the prefrontal cortex.

**Figure 2:**
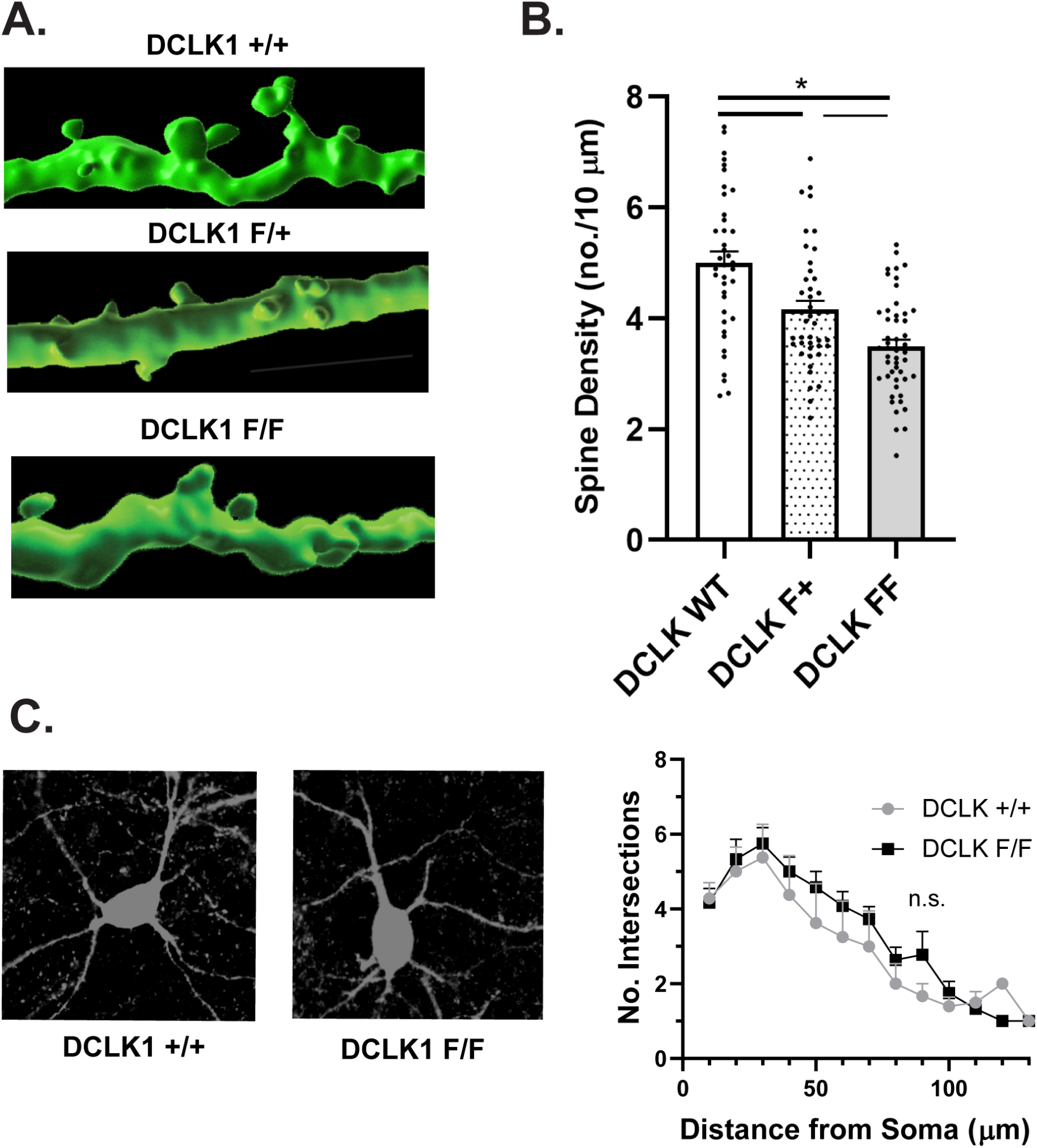
Increased spine density and immature synapses on apical dendrites in DCLK1 F/F pyramidal neurons in PFC layer 2/3. (A) Representative 3D reconstructions of EGFP-labeled apical dendrites and spines on pyramidal neurons in layer 2/3 of prefrontal cortex of DCLK1 +/+, F/+, and F/F mice induced at P10-P13. Scale bar = 10 µm. (B)Quantification of mean spine intensity ± SEM on GFP-labeled apical dendrites of layer 2/3 pyramidal neurons in prefrontal cortex of DCLK1 +/+, F/+, and F/F mice. Each point represents the mean spine density on apical dendrites of a single GFP-expressing pyramidal neuron in 4 DCLK1 +/+ mice, 4 DCLK1 F/+ mice, and 6 DCLK1 F/F mice (P19-34) (*p < 0.05, 2 factor ANOVA with Tukey posthoc comparisons). Approximately 40 neurons/mouse were analyzed. (C)Sholl analysis for branching of dendrites in EGFP-labeled layer 2/3 pyramidal neurons in the prefrontal cortex showed non-significant (n.s.) differences between DCLK1 +/+ and DCLK1 F/F mice (15 neurons/genotype, 3 mice/genotype, P19-P34); paired t-test comparisons of mean number of dendrite crossings at each distance from soma, p< 0.05). Representative images of EGFP-labeled pyramidal neurons of DCLK1 +/+ and DCLK1 F/F mice are shown at the left converted to black and white.

Dendritic spine morphology can be classified based on size and shape (mushroom, stubby, and thin), reflecting the degree of maturity, dynamics, and functional properties (Peters and Kaiserman-Abramof, 1970). Immature spines (stubby and thin) are associated juvenile plasticity and have been termed “learning spines” (Bourne and Harris, 2007), while mushroom spines (“memory spines”) are associated with mature synapses (Bhatt et al., 2009;Holtmaat A. and Svoboda, 2009;Berry and Nedivi, 2017). Deletion of DCLK1 at P10-13 significantly decreased the percent of mature mushroom spines on apical dendrites of layer 2/3 pyramidal neurons in the prefrontal cortex of DCLK1 F/F mice (30 % ± 4) compared to DCLK1 +/+ mice (48% ± 5; *p=0.031, 2-tailed t-test) at P19-P34. There was a corresponding increase in the percent of immature spines (thin and stubby) between DCLK1 F/F (70% ± 3) and DCLK1 +/+ mice (52% ± 6) (p= 0.031) (Fig. 2C).

Sholl analysis was carried out to evaluate any changes in branching of dendritic processes of layer 2/3 pyramidal neurons in the prefrontal cortex (P19-P34) following DCLK1 deletion at P10-P13. No significant differences were observed in branching of DCLK1 +/+ and DCLK1 F/F dendritic arbors at any distance from the soma (paired t-tests for mean number of dendrite crossings at each distance, p < 0.05) (Fig. 2 C,D), indicating that DCLK1 inactivation did not affect dendritic arborization of pyramidal neurons. Representative low magnification images are shown in Fig. 1C. Limitations of dendrite tracing in GFP-expressing neurons in cortical sections precluded full analysis of terminal branching.

These results support a postnatal role for DCLK1 in facilitating spine growth and maturation on apical dendrites of layer 2/3 pyramidal neurons in the prefrontal cortex.

### DCLK1 expression and association with NrCAM at the FIGQY motif

DCLK1 expression was analyzed in the developing postnatal brain by Western blotting of WT mouse cortical lysates during stages of diminished spine remodeling in postnatal mice (P22-P34) and young adults (P50), was well as in older adults (P105) (Fig. 3B). Western blotting for DCLK1 was carried out using an antibody to the carboxyl terminus of full length DCLK1 (82 kDa), which is shared by DCLK1-S (42 kDa). The full length DCLK1 isoform was prominently expressed at all time points in the mouse cortex, whereas DCLK1-S had a very minor presence and was also uniformly expressed (Fig. 3A). To compare the pattern of expression of DCLK1 and NrCAM in cortical lysates in a separate experiment, blots were reprobed for NrCAM (130 kDa). DCLK1 (82 kDa) and NrCAM showed similar patterns of expression at each stage (Fig. 3B). To examine an association of DCLK1 with NrCAM in the developing mouse cortex, NrCAM was immunoprecipitated from equal amounts of cortical lysates followed by immunoblotting for DCLK1. NrCAM and DCLK1 co-immunoprecipitated from cortical lysates at each developmental stage (Fig. 3C). DCLK1-S was not analyzed because it was obscured by the large IgG heavy chain band in the immunoprecipitates. Relative levels of co-immunoprecipitated DCLK1 and NrCAM were generally consistent from P22 to P50, with a small increase in older adults (P105), as indicated by the densitometric ratios of DCLK1/NrCAM (Fig. 3C). The association of DCLK1 with NrCAM was also examined by co-immunoprecipitation from lysates of synaptoneurosomes, a fraction from mouse forebrain (P28) that is enriched in pre-and postsynaptic terminals (Villasana et al., 2006). DCLK1 (82 kDa) also co-immunoprecipitated with NrCAM from synaptoneurosome lysates (Fig. 3D). A nonspecific band of higher molecular weight was evident in immunoprecipitates using either NrCAM antibodies or nonimmune IgG (control). Input blots showed DCLK1 (82 kDa) but not DCLK1-S in the synaptoneurosome fraction.

**Figure 3:**
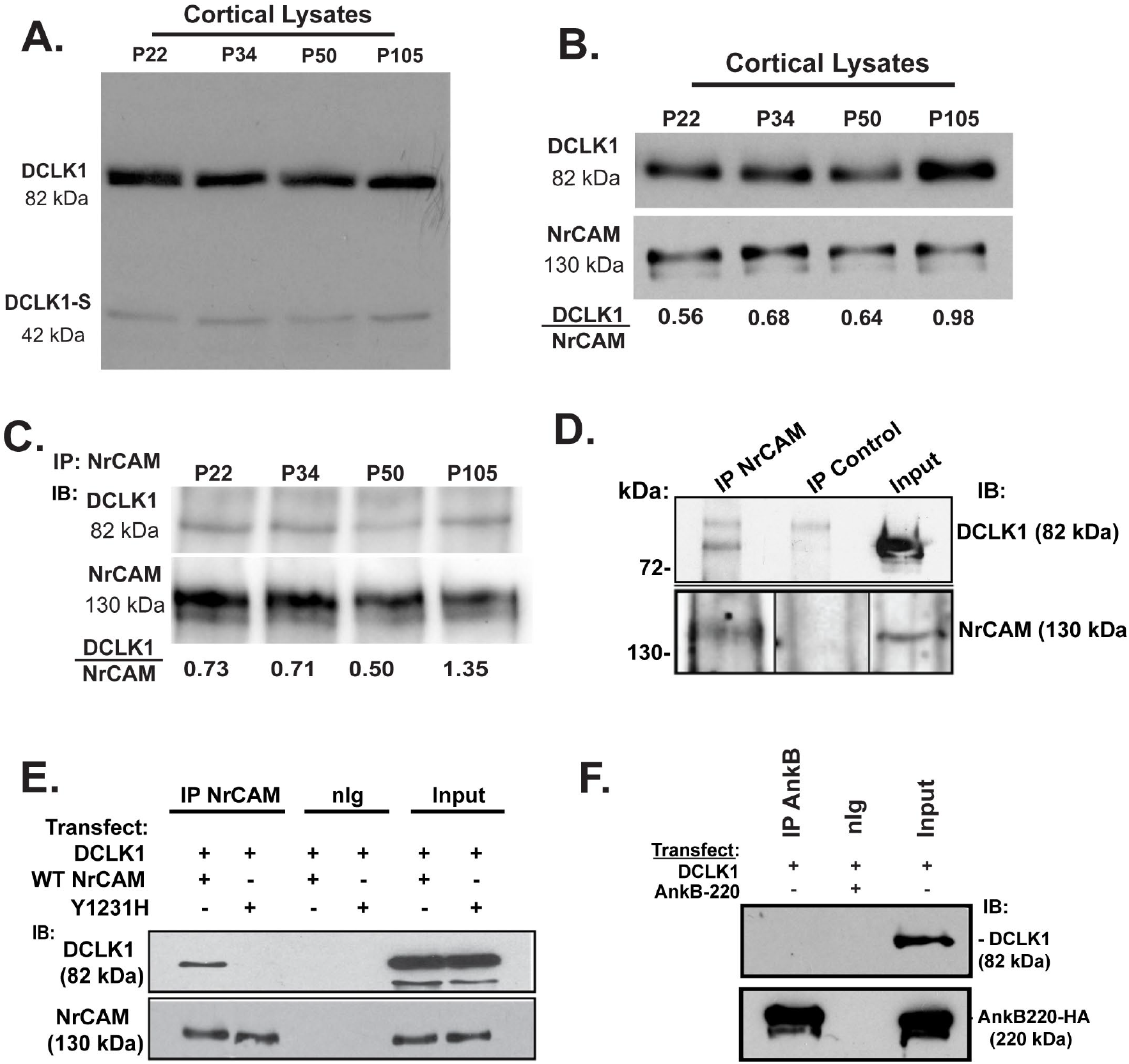
DCLK1 expression and binding to NrCAM at the FIGQY motif. (A) Time course of DCLK1 expression in developing mouse cortex. Western blotting of equal amount of protein lysates from mouse cortex showed the expression of DCLK1 (82 kDa) and DCLK1-S (42kDa) at postnatal stages P22, 34, 50 and older adulthood P105. (B)A different experiment showing expression of DCLK1 (82 kDa) in developing mouse cortical lysates (equal protein) with NrCAM reprobing of the same Western blots (lower panel), indicating coordinate expression from P22-105. Densitometric scanning indicated comparable ratios of DCLK1 to NrCAM in the immunoprecipitates from P22-P50, with an increase at P105. (C)Co-immunoprecipitation of DCLK1 (82 kDa) with NrCAM in mouse cortex lysates at postnatal stages P22, P34, P50 and older adulthood P105. NrCAM was immunoprecipitated (IP) from equal amounts of protein lysates, and immunoblotted (IB) for DCLK1. Blots were reprobed for NrCAM (lower panel). Densitometric scanning indicated comparable ratios of DCLK1 to NrCAM in the immunoprecipitates from P22-P50, with an increase at P105. (D)Co-immunoprecipitation of DCLK1 with NrCAM from mouse brain synaptoneurosomes (P28) (equal protein) shown by immunoprecipitation of NrCAM with NrCAM antibodies or nonimmune IgG (control), followed by immunoblotting (IB) with DCLK1 antibodies. Blots were reprobed for NrCAM (lower panel). Input lysate blots are shown at the right. No DCLK1-S was observed in synaptoneurosome lysates. (E)HEK293T cells were transfected with DCLK, and WT NrCAM or mutant NrCAM Y^1231^H cDNAs, as indicated above. Cell lysates (equal protein) were immunoprecipitated with NrCAM antibodies (IP) or nonimmune IgG, and Western blots probed with DCLK1 antibodies. DCLK1 co-immunoprecipitated with WT NrCAM but not mutant NrCAM Y^1231^H. Blots were reprobed for NrCAM (lower panel). Input lysate blots show comparable levels of expression of DCLK1 and NrCAM in WT and mutant HEK293 cells. (F)HEK293T cells were transfected with AnkB 220-HA and/or DCLK1 cDNAs as indicated above, immunoprecipitated with Ankyrin B antibodies or nonimmune IgG, and immunoblotted for DCLK1. Input lysates are at right. There were no detectable levels of DCLK1 co-immunoprecipitated with Ankyrin B-220 even at prolonged exposures.

The conserved FIGQY motif in the cytoplasmic domain of all L1-CAMs reversibly binds Ankyrins (Bennett and Healy, 2009). Mutation of FIGQY to FIGQH in L1 is a human variant associated with the L1 syndrome of intellectual disability. To test the binding of DCLK1 to NrCAM at this sequence, a tyrosine-1231 to histidine point mutation (Y1231H) was generated in NrCAM cDNA by site directed mutagenesis. HEK293T cells were transfected with expression plasmids for DCLK1 together with WT NrCAM or NrCAM FIGQY^1231^H. Cell lysates were assayed for association of NrCAM and DCLK1 by co-immunoprecipitation. Relative levels of DCLK1 and NrCAM in co-immunoprecipitates were quantified by densitometry. Results showed that DCLK1 co-immunoprecipitated efficiently with WT NrCAM but not with NrCAM FIGQY^1231^H, (Fig. 3E). This finding indicated that DCLK1 interacts directly or indirectly with the NrCAM FIGQY motif. A binding site on DCX for Neurofascin was identified in the second microtubule interaction domain and contains a critical glycine residue for binding (Yap et al., 2018). Mutation of the homologous glycine^259^ to aspartic acid (G259D) in DCLK1 did not abrogate binding to NrCAM in transfected HEK293 cells as determined by coimmunoprecipitation (data not shown).

To determine if DCLK1 associates with NrCAM indirectly through a potential interaction between DCLK1 and Ankyrin B, HEK293T cells were transfected with plasmids expressing DCLK1 and the ubiquitously expressed 220 kDa isoform of Ankyrin B (AnkB-220), then assayed for co-immunoprecipitation. Results revealed no association of DCLK1 with Ankyrin B (Fig.3F), supporting the interpretation that DCLK1 binds to the NrCAM FIGQY motif rather than indirectly through AnkyrinB.

## Discussion

To investigate a role for DCLK1 in the regulation of dendritic spine development in cortical pyramidal neurons, we generated a novel conditional mutant mouse (Nex1Cre-ERT2: DCLK1^*flox/flox*^: RCE) to delete microtubule-binding isoforms of DCLK1, including both full length and DCX-like isoforms, by Cre-mediated recombination of exon 3 under control of the tamoxifen-inducible Nex1 promoter in postmitotic, premigratory pyramidal neurons. Homozygous or heterozygous inactivation of DCLK1 (P10-P13) decreased spine density on apical dendrites of layer 2/3 pyramidal neurons in the prefrontal cortex. Loss of DCLK1 preferentially decreased the percent of mature (mushroom) spines, and increased the percent of immature spines (thin and stubby). DCLK1 was expresssed coordinately with neural cell adhesion molecule NrCAM across postnatal and adult development. DCLK1 was found to bind NrCAM at a conserved sequence FIGQY in its cytoplasmic domain, which is known to recruit the spectrin-actin adaptor Ankyrin. These results support a role for DCLK1 in facilitating spine growth and maturation during postnatal development of the prefrontal cortex.

The phenotype of DCLK1 F/F mice shares similarities and differences with other DCLK1 mouse models. Induction of recombination in Nex1Cre-ERT2: DCLK1 F/F mice at P10-P13 resulted in decreased spine density on apical dendrites of cortical pyramidal neurons but no obvious alteration in cortical lamination, dendritic branching, or major axon tracts. Similarly, a global DCLK1 knockout mouse derived via homologous recombination in embryonic stem cells exhibited normal survival, gross brain architecture, and migration of neuronal progenitors (Deuel et al., 2006). Excision of DCLK1 exon 3 in the EIIa-Cre: DCLK1 conditional mutant causes germline recombination and removes isoforms with microtubule-binding domains leaving the kinase-only isoform intact (Koizumi et al., 2006). The EIIa-Cre mutant showed defects in formation of the corpus callosum and anterior commissure, unlike our mutant, in which excision was onduced later at postnatal stages. In a different approach (Shin et al., 2013), DCLK1 expression was downregulated by *in utero* shRNA expression in neuronal progenitors at embryonic day E15.5, resulting in suppression of dendritic growth and branching in the somatosensory cortex. Slight increases in spine density and spine width were also noted. In our Nex1Cre-ERT2 mouse model postnatal deletion of DCLK1 isoforms containing microtubule binding domains resulted in a significant decrease in spine density, preferentially targeting mature spines with mushroom morphology, and did not decrease dendritic arborization detectably. Such differences may be due to the later time frame of DCLK1 recombination in Nex1Cre-ERT2: DCLK1^flox/flox^ : RCE mice compared to EIIa-induced Cre recombination or shRNA expression, both of which occur in immature neural progenitors. One possibility is that DCLK1 in immature neurons may preferentially promote dendritic growth, building the dendritic tree through microtubule transport of proteins for dendrite extension and arborization. Later, in differentiating pyramidal neurons, DCLK1 may promote spine growth and maturation using microtubules to transport proteins into developing spines via direct deposit or membrane diffusion (Dent, 2017). This scenario is in accord with the ability of DCLK1 to promote dendritic branching in neuronal cultures at early (DIV4) but not later (DIV9) time points (Shin et al., 2013).

In the present study we show that DCLK1 binds to NrCAM at postnatal stages of development in the mouse neocortex, and this binding is dependent on the FIGQY motif within the NrCAM cytoplasmic domain. The lissencephaly protein DCX, which shares sequence homology with DCLK1 in the microtubule binding domains, was reported to bind the L1-CAM Neurofascin at the conserved FIGQY motif, but not NrCAM or L1, indicating specificity of association between L1-CAMs and DCX family members (Kizhatil et al., 2002). The FIGQY motif is the location at which Ankyrin reversibly engages L1-CAMs (Bennett and Healy, 2009). Tyrosine phosphorylation of the FIGQY motif in Neurofascin decreases the affinity for Ankyrin allowing recruitment of DCX (Kizhatil et al., 2002). DCLK1 did not co-immunoprecipitate with Ankyrin B from transfected HEK293 cells, suggesting that DCLK1 may associate directly with NrCAM at this sequence.

DCLK1 might serve to promote or maintain spine growth and maturation during spine morphogenesis through its ability to transport dense core vesicles along microtubules (Lipka et al., 2016). For example, DCLK1 may bind and transport NrCAM to the spine plasma membrane, positioning it for intercellular adhesion or spine pruning in response to activity-dependent expression of Sema3F (Mohan et al., 2019a). After spines are produced, there may be a temporally regulated shift of NrCAM binding from DCLK1 to Ankyrin, possibly regulated by tyrosine dephosphorylation of NrCAM at the FIGQY motif. With respect to additional modes of DCLK1 regulation, recent research has shown that DCLK1 autophosphorylates on its carboxyl terminal tail, which inhibits microtubule binding (Agulto et al., 2021). Future studies with the Nex1Cre-ERT2: DCLK1^flox/flox^ : RCE mice will be aimed at determining if loss of DCLK1 from pyramidal neurons impairs excitatory transmission or behavior due to reduction of spine density on apical dendrites of cortical pyramidal neurons in the prefrontal cortex. In addition, recent studies demonstrated that DCLK1 knockdown by shRNA in a mouse model of Parkinson’s disease-like synucleopathy mitigates dopaminergic neurotoxicity and downregulates α-Synuclein levels posttranscriptionally (Vazquez-Velez et al., 2020). Since DCLK1 is a druggable target, Nex1Cre-ERT2: DCLK1^flox/flox^: RCE mice could be useful for investigating deleterious consequences of adult knockout of DCLK1 in pyramidal neurons.

## Abbreviations

(DCLK1): Doublecortin-like kinase-1
(DCX): Doublecortin
(NrCAM): Neuron-glia related cell adhesion molecule
(Sema3): Semaphorin 3

## Acknowledgements

This work was supported by the US National Institutes of Mental Health grant R01 MH113280 (PFM), UNC School of Medicine Biomedical Research Core Project award (PFM), Carolina Institute for Developmental Disabilities center grant NIH P50HD103573 (Dr. Joseph Piven, PI), and T32 NRSA (5T32HD040127-18)(KEM). We acknowledge Dr. Pablo Ariel, Director of the Microscopy Services Laboratory in the UNC Department of Pathology and Laboratory Medicine, who provided expert advice on imaging (P30 CA016086 Cancer Center Core Grant). We thank Dr. Gordon Fishell for providing the RCE mouse strain, and Alex Kampov-Polevoi for technical assistance.

